# Meta-transcriptomic identification of divergent *Amnoonviridae* in fish

**DOI:** 10.1101/2020.10.06.329003

**Authors:** Olivia M. H. Turnbull, Ayda Susana Ortiz-Baez, John-Sebastian Eden, Mang Shi, Jane E. Williamson, Troy F. Gaston, Yong-Zhen Zhang, Edward C. Holmes, Jemma L. Geoghegan

## Abstract

Tilapia lake virus (TiLV) has caused mass mortalities in farmed and wild tilapia with serious economic and ecological consequences. Until recently, this virus was the sole member of the *Amnoonviridae*, a family within the order *Articulavirales* comprising segmented negative-sense RNA viruses. We sought to identify additional viruses within the *Amnoonviridae* through total RNA sequencing (meta-transcriptomics) and data mining of published transcriptomes. Accordingly, we sampled marine fish species from both Australia and China and discovered two new viruses within the *Amnoonviridae*, tentatively called *Flavolineata virus* and *Piscibus virus*, respectively. In addition, by mining vertebrate transcriptome data we identified nine additional virus transcripts matching to multiple genomic segments of TiLV in both marine and freshwater fish. These new viruses retained sequence conservation with the distantly related *Orthomyxoviridae* in the RdRp subunit PB1, but formed a distinct and diverse phylogenetic group. These data suggest that the *Amnoonviridae* have a broad host range within fish and that greater animal sampling will identify additional divergent members of the *Articulavirales*.

## 1. Introduction

The *Amnoonviridae* are a recently described family of segmented and enveloped negative-sense RNA viruses associated with disease in fish. Until recently, the *Amnoonviridae* comprised only a single species, *tilapinevirus* or tilapia lake virus (TiLV) [1, 2], that is associated with high rates of morbidity and mortality in both farmed and wild tilapia (*Oreochromis niloticus* and *Oreochromis niloticus* x *O. aureus* hybrid). As the second most farmed fish globally [3] and an important subsistence for farmers and high value markets [4], tilapia contribute $7.5 billion annually to the aquaculture industry. Outbreaks of TiLV have resulted in significant economic and ecological loss. The virus causes gross lesions of the eyes and skin, while also impacting brain, liver and kidney tissue [2], with associated mortality rates up to 90% [5, 6]. While ongoing surveillance has detected the virus across numerous countries in Asia, Africa and South America [7], the identity of potential reservoir hosts remains unclear.

The *Amnoonviridae* are members of the order *Articulavirales* that also includes the *Orthomyxoviridae* [8] that are particularly well-known because they contain the mammalian and avian influenza viruses. Unlike the *Amnoonviridae*, the *Orthomyxoviridae*, and closely related but unclassified orthomyxo-like viruses, infect a broad range of host species comprising both invertebrates and vertebrates. Notably, a divergent member of the *Amnoonviridae*, *Lauta virus*, was recently identified in an Australian gecko [9], strongly suggesting that members of this family are present in a wider range of vertebrate hosts. In addition, the large phylogenetic distance between *Lauta virus* and TiLV suggests that the former may even constitute a new genus within the *Amnoonviridae*, with the long branches throughout the *Articulavirales* phylogeny likely indicative of very limited sampling.

To help address whether the *Amnoonviridae* might be present in a wider range of vertebrate taxa we screened for their presence using a meta-transcriptomic analysis of marine fish sampled in Australia and China, combined with transcriptome mining.

## 2. Materials and Methods

### 2.1 Fish collection in Australia

Fish samples were collected from Bass Strait (40°15′S–42°20′S, 147°05′E–148°35′E), Australia in November 2018. The fish species collected included *Rhombosolea tapirina, Platycephalus bassensis, Platycephalus speculator, Trachurus declivis, Trachurus novaezelandiae, Scorpaena papillosa, Pristiophorus nudipinnis, Pentaceropsis recurvirostris* and *Meuschenia flavolineata.* Fish were caught via repeated research trawls on the fisheries training vessel, *Bluefin*, following the methodology outlined in [10]. Ten individuals from each species were caught and stored separately. Gill tissues were dissected and snap frozen at −20°C on the vessel, and then stored in a −80°C freezer at Macquarie University, Sydney. Sampling was conducted under the approval of the University of Tasmania Animal Ethics Committee, approval number A0015366

### 2.2 Fish collection in China

Fish samples were collected from South China Sea as reported previously [11]. Fish species that were sampled and subsequently pooled included *Proscyllium habereri*, *Urolophus aurantiacus*, *Rajidae sp.*, *Eptatretus burgeri*, *Heterodontus zebra*, *Dasyatis bennetti, Acanthopagrus latus, Epinephelus awoara, Conger japonicus, Siganus canaliculatus, Glossogobius circumspectus, Halichoeres nigrescens*, and *Boleophthalmus pectinirostris*. Liver samples from each species were pooled and stored in a −80°C freezer. The procedures for sampling and sample processing were approved by the ethics committee of the National Institute for Communicable Disease Control and Prevention of the China CDC.

### 2.3 RNA sequencing

For RNA extraction, frozen tissue was partially thawed and submerged in lysis buffer containing 1% ß-mercaptoethanol and 0.5% Reagent DX before tissues were homogenized together with TissueRupture (Qiagen). The homogenate was centrifuged to remove any potential tissue residues, and RNA from the clear supernatant was extracted using the Qiagen RNeasy Plus Mini Kit. RNA was quantified using NanoDrop (ThermoFisher). RNA isolated from the Australian samples was pooled for each host species, whereas RNA isolated from the Chinese samples was pooled from all species, resulting in 3μg per pool (250ng per individual). Libraries were constructed using the TruSeq Total RNA Library Preparation Protocol (Illumina) and host ribosomal RNA (rRNA) was depleted using the Ribo-Zero-Gold Kit (Illumina) to facilitate virus discovery. Fish caught in Australia were subject to paired-end (100 bp) sequencing performed on the NovaSeq 500 platform (Illumina) carried out by the Australian Genome Research Facility (AGRF). RNA sequencing of the pooled fish sampled from China were sequenced on the HiSeq 2500 platform (Illumina) at BGI Tech (Shenzhen).

### 2.4 Transcript sequence similarity searching for novel amnoonviruses

Sequencing reads were first quality trimmed then assembled *de novo* using Trinity RNA-Seq (v.2.11.0) [12]. The assembled contigs were annotated based on similarity searches against the National Center for Biotechnology Information (NCBI) nucleotide (nt) and non-redundant protein (nr) databases using BLASTn and Diamond BLASTX (v.2.0.2) [13]. To infer the evolutionary relationships of the amnoonviruses newly discovered the translated viral contigs were combined with representative protein sequences from TiLV and *Lauta virus* obtained from NCBI GenBank. The sequences retrieved were then aligned with those generated here using MAFFT (v7.4) employing the E-INS-i algorithm. Ambiguously aligned regions were removed using trimAl (v.1.2) [14]. To estimate phylogenetic trees, we utilized the maximum likelihood approach available in IQ-TREE (v 1.6.8) [15], selecting the best-fit model of amino acid substitution with ModelFinder [16], and using 1000 bootstrap replicates. Phylogenetic trees were annotated with FigTree (v.1.4.2).

### 2.5 PCR confirmation

To further confirm the presence of *Flavolineata virus* in the yellow-striped leatherjacket collection, 10µl of extracted RNA was transcribed into cDNA using SuperScript® VILO™ reverse transcriptase (Invitrogen, CA USA). PCR amplification was performed using Platinum™ II Hot-Start PCR Master Mix (2X) (Invitrogen, CA, USA) and 3 sets of primers (Table S1) designed to cover different regions of the virus sequence. PCR products were visualized on 2% agarose gel stained with SYBR® Safe (Invitrogen, CA USA).

### 2.6 TSA mining

To identify additional novel vertebrate viruses within the *Amnoonviridae* we screened *de novo* transcriptome assemblies available at the NCBI Transcriptome Shotgun Assembly (TSA) database (https://www.ncbi.nlm.nih.gov/genbank/tsa/). Amino acid sequences of *Flavolineata virus*, *Piscibus virus* and TiLV were queried against the assemblies using the translated Basic Local Alignment Search Tool (tBLASTn) algorithm. We restricted the search to transcriptomes within the Vertebrata (taxonomic identifier: 7742). Putative virus contigs were subsequently queried using BLASTx against the non-redundant virus database.

### 2.7 Virus naming

New viruses identified in this study are tentatively named by drawing from their host species’ names.

### 2.8 Data availability

Sequencing reads are available at the NCBI Sequence Read Archive (SRA). For *Piscibus virus* see Bioproject PRJNA418053 (BioSample: SAMN08013970; Library name: BHFishG) and for *Flavolineata virus* see Bioproject: PRJNA667570. Alignments with new virus transcripts are available at https://github.com/jemmageoghegan/Amnoonviridae-in-fish.

## 3. Results

### 3.1 Identification of a novel Amnoonviridae in yellow-striped leatherjacket

As part of a large virological survey on nine species of marine fish our meta-transcriptomic analysis identified a novel member of the *Amnoonviridae*, tentatively named *Flavolineata virus*, in a sequencing library of 10 pooled individuals of yellow-striped leatherjacket (*Meuschenia flavolineata*) sampled from the Bass Strait off the coast of Tasmania, Australia. No amnoonviruses were identified in the remaining eight fish species. We identified a complete, highly divergent protein in which a Diamond BLASTx analysis revealed 37% amino acid identity to TiLV segment 1, characterized as the PB1 subunit (Genbank accession: QJD15207.1, e-value: 2.0×10-80, query coverage 95%), with a GC composition of 48.2% and a standardised abundance of 0.00004% of the total non-rRNA library. The presence of *Flavolineata virus* was further confirmed in the unpooled samples using RT-PCR (Figure S1).

### 3.2 Identification of a novel Amnoonviridae in pooled marine fish from the South China Sea

An additional novel member of the *Amnoonviridae*, in which we have termed *Piscibus virus*, was identified in a pool of various marine species (including sharks, eels, stingrays, jawless fish and perch-like fish) sampled in the South China Sea as described previously [11]. Specifically, we identified a short contig (270 nucleotides) that shared highest amino acid sequence similarity (48.8%, e-value: 1.5×10^−15^) to *Flavolineata virus* using a custom database including the known members of the family *Amnoonviridae*. In addition, a comparison to the NCBI nr database showed that *Piscibus virus* had 48.5% amino acid similarity (e-value: 2.0×10^−07^) to the PB1 subunit of the TiLV RdRp. The GC composition of the assembled sequence was 49.2% and it had a standardized abundance of 0.0001% of the total non-rRNA library. Despite the limited contig length, we identified conserved motifs within the PB1 subunit (see below).

### 3.3 Identification of novel Amnoonviridae in published transcriptomes

To identify additional novel vertebrate viruses within the *Amnoonviridae* we screened *de novo* transcriptome assemblies available at NCBI’s TSA database. In doing so we identified nine further potentially novel viruses in fish (Table 1).

**Table 1.** Novel viruses identified in this study.

| **Virus name** | **Host** | **Detection method (NCBI accession of TSA data)** | **Contig length matching segment 1 of TiLV (nt)** | **Closest amino acid match to segment 1 (GenBank accession)** |
| --- | --- | --- | --- | --- |
| *Flavolineata virus* | *Meuschenia flavolineata* | Fish sampling + meta-transcriptomics | 1536 | 37% TiLV (QJD15207.1) |
| *Piscibus virus* | Pooled marine fish (see methods) | Fish sampling + meta-transcriptomics | 270 | 49% TiLV (QJD15207.1) |
| *Dolomieu virus* | *Micropterus dolomieu* | TSA search<br>(GDQU01066121.1,<br>GDQU01106321.1,<br>GDQU01283605.1,<br>GDQU01532168.1) | 1440 | 34% TiLV (QJD15204.1) |
| *Namensis virus* | *Gymnocypris namensis* | TSA search<br>(GHYH01080462.1,<br>GHYH01005036.1,<br>GHYH01084204.1) | 1503 | 35% TiLV (AOE22913.1) |
| *Hamatus virus* | *Chionodraco hamatus* | TSA search<br>(GFMN01088333.1) | 321 | 51% TiLV (QJD15205.1) |
| *Stewartii virus* | *Oxygymnocypris stewartii* | TSA search<br>(GIBO01031171.1,<br>GIBO01013027.1) | 1743 | 28% TiLV (QJD15204.1) |
| *Plagiostomus virus* | *Schizothorax plagiostomus* | TSA search<br>(GHXZ01024367.1,<br>GHXZ01079240.1) | 366 | 39% TiLV (AOE22912.1) |
| *Przewalskii virus* | *Gymnocypris przewalskii* | TSA search<br>(GHYJ01002273.1,<br>GHYJ01008047.1,<br>GHYJ01010906.1) | 1761 | 26% TiLV (QJD15208.1) |
| *Asotus virus 1* | *Silurus asotus* | TSA search<br>(GHGF01026383.1,<br>GHGF01034639.1,<br>GHGF01033499.1,<br>GHGF01028660.1,<br>GHGF01037407.1) | 1710 | 32% TiLV (QMT29723.1) |
| *Asotus virus 2* | *Silurus asotus* | TSA search<br>(GHGF01016319.1,<br>GHGF01027066.1,<br>GHGF01047620.1) | 1719 | 29% TiLV (QJD15204.1) |
| *Nudifrons virus* | *Lindbergichthys nudifrons* | HACN01008153.1 | 1032 (segment 4) | 44% *Flavolineata virus* (segment 4) |

Relatives of the *Amnoonviridae* were identified in ray-finned fish species (Actinopterygii) from marine (*Lepidonotothen nudifrons* and *Chionodraco hamatus*) and freshwater ecosystems (*Gymnocypris przewalskii*, *Gymnocypris namensis, Micropterus dolomieu, Oxygymnocypris stewartia*, *Schizothorax plagiostomus* and *Silurus asotus*) (Table 1). All viral sequences corresponded to segments 1-4 and ranged from 209-1784 nucleotides in length. The putative segments shared 26-51% sequence identity with TiLV. Most of the identified viral sequences corresponded to segment 1, containing the RdRp and covered motifs II and III (Figure 2). Notably, no other vertebrate class within the TSA were identified as potential hosts of these viruses.

### 3.4 Evolutionary relationships of novel Amnoonviridae

We next performed phylogenetic analysis of the RdRp subunit (segment 1) across the order *Articulavirales* (Figure 1). This revealed two distinct clades of fish viruses within the *Amnoonviridae* (with 83% bootstrap support). The original member of this virus family, TiLV, grouped with *Flavolineata virus*, *Piscibus virus*, *Dolomieu virus*, *Namensis virus* and *Hamatus virus* in one clade. The second fish virus clade comprised the newly identified *Stewartii virus*, *Plagiostomus virus*, *Przewalskii virus*, *Asotus virus 1*, and *Asotus virus 2*. *Lauta virus*, identified in a native Australian gecko, appears to form a distinct lineage, suggestive of a separate genus. This phylogenetic analysis clearly illustrates the diversity of these viruses within both marine and freshwater fish, with no apparent host taxonomic structure (Figure 1).

**Figure 1.**
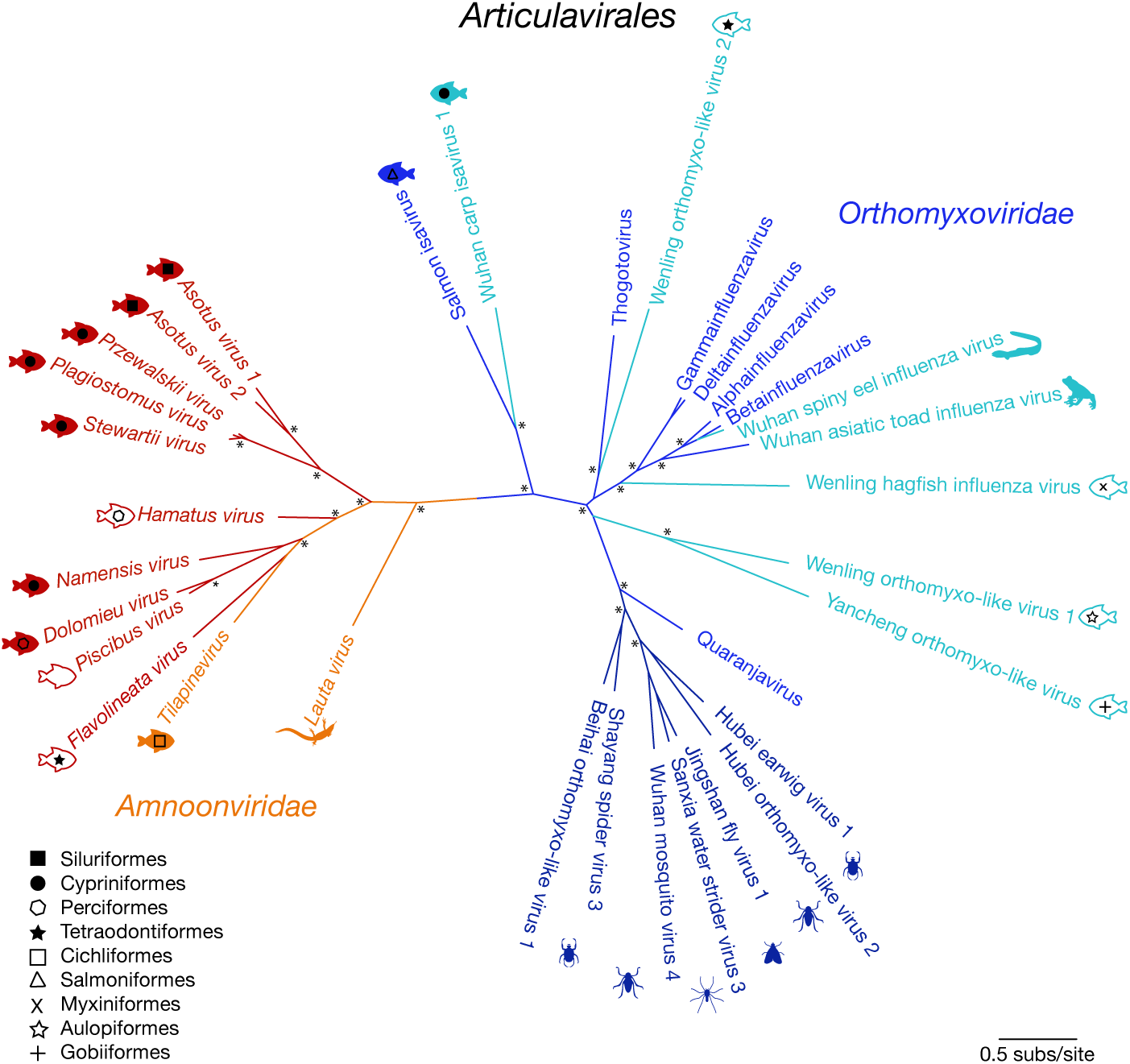
Unrooted maximum likelihood phylogenetic tree of the PB1 subunit showing the topological position of ten of the 11 newly discovered viruses (red) that shared sequence similarity to segment 1 within the order *Articulavirales* (*Amnoonviridae*: orange; *Orthomyxoviridae*: shades of blue). Tilapinevirus (TiLV) and the recently discovered *Lauta virus* were the only viruses previously identified in this family. Fish viruses are annotated with fish symbols (filled: freshwater; outline: marine) and fish order corresponds to shapes illustrated by the key. All branches are scaled according to the number of amino acid substitutions per site. An asterisk (*) illustrates nodes with bootstrap support >70%.

### 3.5 Genome composition of the novel Amnoonviridae

Ten of the novel viruses identified included segment 1, corresponding to the RdRp subunit PB1, and sharing clear sequence homology with different members of the *Articulavirales* including the *Orthomyxoviridae* (Figure 2). These viruses had a closest genetic match to TiLV, ranging from 28-51% sequence similarity at the amino acid level to segment 1 (Table 1). Segment 4 was the only segment found from the tentatively named nudifrons virus (Table 1).

**Figure 2.**
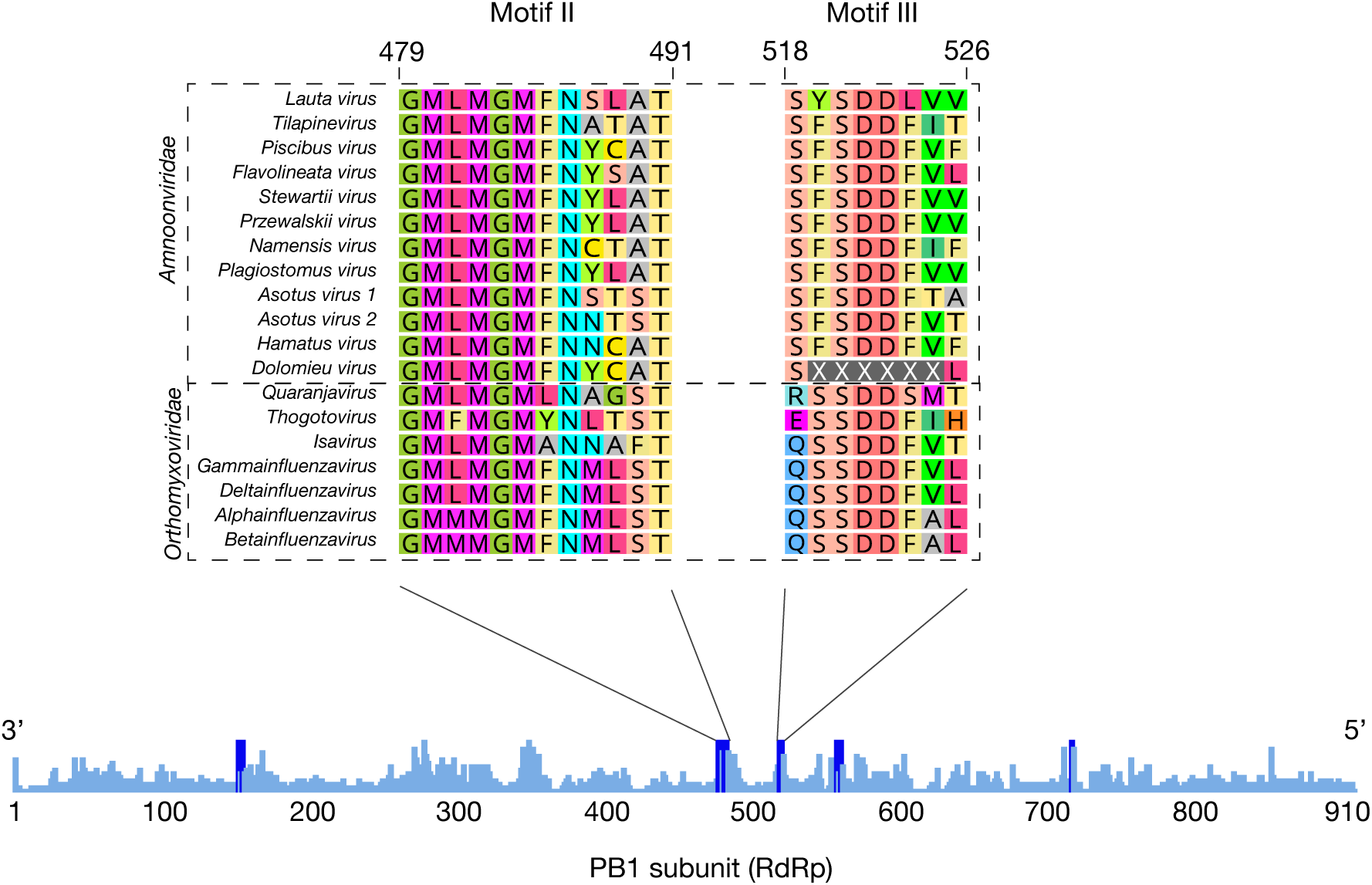
Alignment of viruses within the order *Articulavirales*, highlighting conserved motifs in the RdRp subunit PB1. The sequences of two motifs (II and III) are shown where all sequences overlapped.

Despite the lack of genomic characterization of TiLV, a sequence comparison across the *Articulavirales* revealed conserved PB1 motifs (Figure 2). Sequence similarities with other viral RNA-dependent RNA polymerases suggest that motif III plays a key functional role at the core of the transcriptase-replicase activity [17, 18]. Defined by the consensus serine-aspartic acid-aspartic acid (SDD) sequence in the *Articulavirales*, this motif is highly conserved and is critical for protein stability and function.

In contrast to the six to eight genomic coding segments that comprise viruses within the *Orthomyxoviridae*, TiLV contains 10 segments with open reading frames, of which only segment 1 has been functionally characterized to date [19]. While we were able to distinguish virus transcripts with sequence similarity to segments 1 – 4 of TiLV (Figure 3), it is possible that the other segments are present but too divergent in sequence to be detected. Indeed, the remaining segments of TiLV exhibit no sequence similarity to any other known viruses [1, 2] or eukaryotic genes.

**Figure 3.**
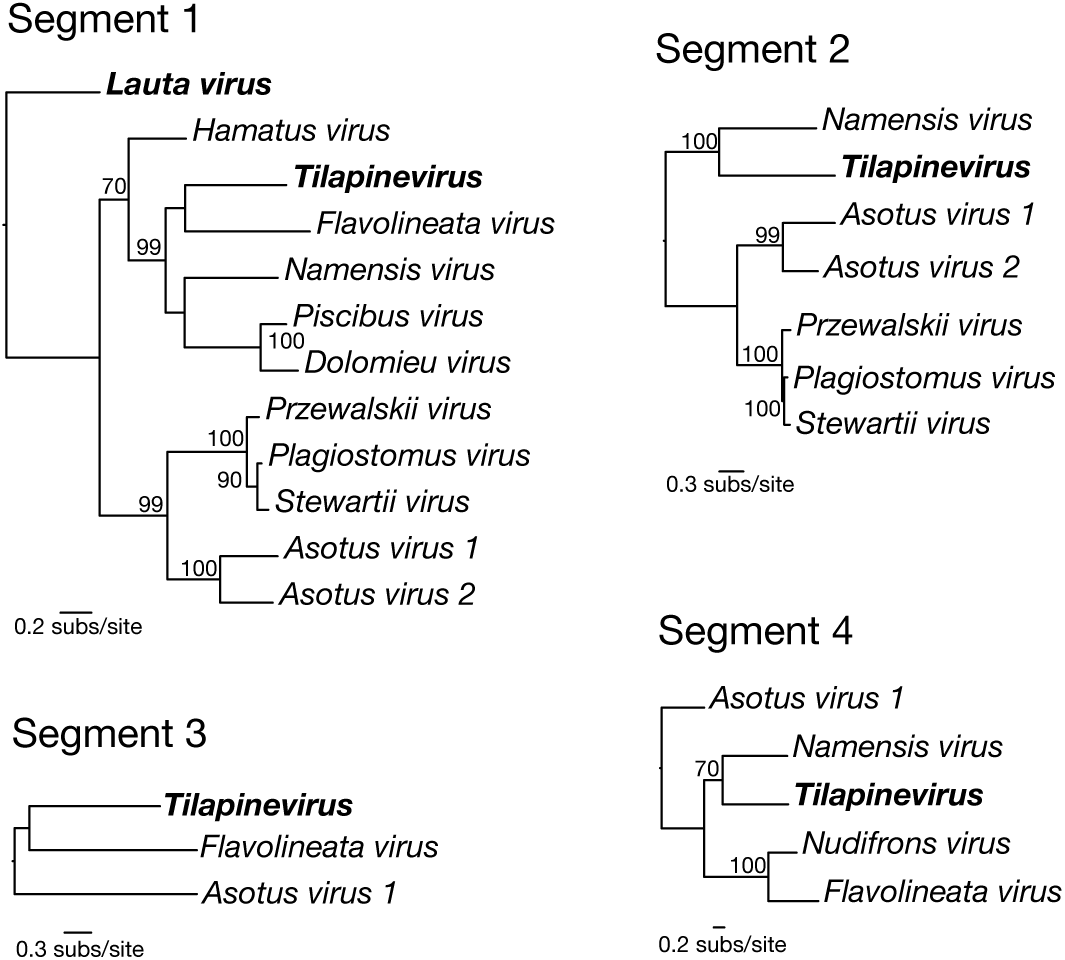
Maximum likelihood phylogenetic trees of genomic segments 1-4 for the new virus transcripts identified in this study within the *Amnoonviridae*. Viruses previously identified in this family are in bold. Bootstrap values >70% are shown. The segment 1 phylogeny was rooted using *Lauta virus* as the outgroup (as suggested by the tree in Figure 1). The remaining three segment phylogenies were then rooted to match the segment 1 tree. Branch scale bars are shown for each, representing the number of substitutions per site.

Also of note was that we found some evidence for phylogenetic incongruence between the topologies of the different gene segments, although this analysis is complicated by the differing numbers of viruses available for each segment, the short sequence alignments, and the highly divergent nature of the sequences being analysis. For example, *Flavolineata virus* and TiLV appear as sister taxa in segment 1 yet are seemingly more divergent in segment 4 (Figure 3). Hence, this phylogenetic pattern tentatively suggests that amnoonviruses may have undergone reassortment in similar manner to influenza A viruses in the *Orthomyxoviridae*, although this will need to be confirmed with the addition of longer sequences and more taxa. Reassortment has previously been observed within circulating TiLV strains, which has added complexity to inferring its evolutionary history [20].

## 4. Discussion

Through both sampling marine fish and mining publicly available sequence data, we discovered 11 new viruses, all of which are the closest genetic relatives of TiLV. These viruses fall within the *Amnoonviridae*, which currently comprises only two viruses: TiLV and *Lauta virus*. The discovery of these new viruses expands our understanding of the host range of the *Amnoonviridae* to include host species across multiple taxonomic orders of freshwater and marine fish, including Cypriniformes, Siluriformes, Perciformes and Tetraodontiformes, and includes animals sampled in a range of geographic localities (Australia, China, North America, Antarctica and Japan). Not only does the identification of these new viruses greatly increase the phylogenetic diversity in this newly identified group of viruses, but it may also provide insight into the potential origins and host range of TiLV, a virus that has major economic and ecological impacts on fisheries and aquaculture.

The viruses discovered here were highly divergent in sequence, likely limiting our ability to detect all genome segments present in the data. Nevertheless, sequence conservation within segment 1 across the entire taxonomic order strongly supports the inclusion of these new viruses within the *Amnoonviridae*. While we only found new viruses in fish and no other vertebrate classes, it is important to note that fish comprise 44% of currently available vertebrate transcriptomes (as of September 2020). With the expansion of these databases, it is likely we will identify additional highly divergent viruses within the *Amnoonviridae* and hence of the *Articulavirales* as a whole. The discovery of these 11 viruses invites further research into the true diversity and evolutionary origins of the *Amnoonviridae*.

## Supplementary Materials

The following are available online at www.mdpi.com/xxx/s1: Table S1: List of primer sets used for the RT-PCR confirmation of *Flavolineata virus* in specimens of *Meuschenia flavolineata*; Table S2: All virus transcripts identified in this study that fell across genomic segments within the *Amnoonviridae*; Figure S1: Agarose gels electrophoresis showing PCR products from three sets of primers that target a region in the PB1 gene segment (RdRp) for ten individuals of *Meuschenia flavolineata*.

## Author Contributions

Conceptualization, E.C.H. and J.L.G.; formal analysis, O.M.H.T, A.S.O.B. J.S.E, J.L.G; resources, J.E.W., T.F.G., M.S., Y.Z.Z, E.C.H and J.L.G.; writing—original draft preparation, O.M.H.T, A.S.O.B., J.S.E., E.C.H. and J.L.G.; writing—review and editing, all authors; funding acquisition, E.C.H. and J.L.G. All authors have read and agreed to the published version of the manuscript.

## References

1. Bacharach, E.; Mishra, N.; Briese, T.; Zody, M. C.; Kembou Tsofack, J. E.; Zamostiano, R.; Berkowitz, A.; Ng, J.; Nitido, A.; Corvelo, A.; Toussaint, N. C.; Abel Nielsen, S. C.; Hornig, M.; Del Pozo, J.; Bloom, T.; Ferguson, H.; Eldar, A.; Lipkin, W. I., Characterization of a novel orthomyxo-like virus causing mass die-offs of tilapia. mBio 2016, 7, (2), e00431–16.

2. Eyngor, M.; Zamostiano, R.; Kembou Tsofack, J. E.; Berkowitz, A.; Bercovier, H.; Tinman, S.; Lev, M.; Hurvitz, A.; Galeotti, M.; Bacharach, E.; Eldar, A., Identification of a novel RNA virus lethal to tilapia. J Clin Microbiol 2014, 52, (12), 4137–4146.

3. Fishery and aquaculture statistics; http://www.fao.org/3/i3507t/i3507t.pdf (accessed Sept 2020) 2011.

4. Fitzsimmons, K., Global tilapia market update 2015. DOI: 10.13140/RG.2.1.2848.9448 2015.

5. Behera, B. K.; Pradhan, P. K.; Swaminathan, T. R.; Sood, N.; Paria, P.; Das, A.; Verma, D. K.; Kumar, R.; Yadav, M. K.; Dev, A. K.; Parida, P. K.; Das, B. K.; Lal, K. K.; Jena, J. K., Emergence of tilapia lake virus associated with mortalities of farmed Nile tilapia *Oreochromis niloticus* (Linnaeus 1758) in India. Aquaculture 2018, 484, 168–174.

6. Surachetpong, W.; Janetanakit, T.; Nonthabenjawan, N.; Tattiyapong, P.; Sirikanchana, K.; Amonsin, A., Outbreaks of tilapia lake virus infection, Thailand, 2015-2016. Emerg Infect Dis 2017, 23, (6), 1031–1033.

7. Jansen, M. D.; Dong, H. T.; Mohan, C. V., Tilapia lake virus: a threat to the global tilapia industry? Rev Aquac 2019, >11, (3), 725–739.

8. Virus Taxonomy: 2018a Release. https://ictv.global/taxonomy

9. Ortiz-Baez, A. S.; Eden, J.-S.; Moritz, C.; Holmes, E. C., A divergent articulavirus in an Australian gecko identified using meta-transcriptomics and protein structure comparisons. Viruses 2020, 12.

10. Park, J. M.; Coburn, E.; Platell, M. E.; Gaston, T. F.; Taylor, M. D.; Williamson, J. E., Diets and resource partitioning among three sympatric gurnards in northeastern Tasmanian waters, Australia. Mar Coast Fish 2017, 9, (1), 305–319.

11. Shi, M.; Lin, X.-D.; Chen, X.; Tian, J.-H.; Chen, L.-J.; Li, K.; Wang, W.; Eden, J.-S.; Shen, J.-J.; Liu, L.; Holmes, E. C.; Zhang, Y. Z., The evolutionary history of vertebrate RNA viruses. Nature 2018, 556, 197–202.

12. Haas, B. J.; Papanicolaou, A.; Yassour, M.; Grabherr, M.; Blood, P. D.; Bowden, J.; Couger, M. B.; Eccles, D.; Li, B.; Lieber, M.; MacManes, M. D.; Ott, M.; Orvis, J.; Pochet, N.; Strozzi, F.; Weeks, N.; Westerman, R.; William, T.; Dewey, C. N.; Henschel, R.; LeDuc, R. D.; Friedman, N.; Regev, A., *De novo* transcript sequence reconstruction from RNA-Seq: reference generation and analysis with Trinity. Nat Protoc 2013, 8, (8), 10.1038/nprot.2013.084.

13. Buchfink, B.; Xie, C.; Huson, D. H., Fast and sensitive protein alignment using DIAMOND. Nat Methods 2015, 12, (1), 59–60.

14. Capella-Gutierrez, S.; Silla-Martinez, J. M.; Gabaldon, T., trimAl: a tool for automated alignment trimming in large-scale phylogenetic analyses. Bioinformatics 2009, 25, (15), 1972–3.

15. Nguyen, L.-T.; Schmidt, H. A.; von Haeseler, A.; Minh, B. Q., IQ-TREE: a fast and effective stochastic algorithm for estimating maximum-likelihood phylogenies. Mol Biol Evol 2015, 32, (1), 268–274.

16. Kalyaanamoorthy, S.; Minh, B. Q.; Wong, T. K. F.; von Haeseler, A.; Jermiin, L. S., ModelFinder: fast model selection for accurate phylogenetic estimates. Nat Methods 2017, 14, (6), 587–589.

17. Chu, C.; Fan, S.; Li, C.; Macken, C.; Kim, J. H.; Hatta, M.; Neumann, G.; Kawaoka, Y., Functional analysis of conserved motifs in influenza virus PB1 protein. PLoS One 2012, 7, (5), e36113–e36113.

18. Biswas, S. K.; Nayak, D. P., Mutational analysis of the conserved motifs of influenza A virus polymerase basic protein 1. J Virol 1994, 68, (3), 1819–26.

19. Taengphu, S.; Sangsuriya, P.; Phiwsaiya, K.; Debnath, P. P.; Delamare-Deboutteville, J.; Mohan, C. V.; Dong, H. T.; Senapin, S., Genetic diversity of tilapia lake virus genome segment 1 from 2011 to 2019> and a newly validated semi-nested RT-PCR method. Aquaculture 2020, 526, 735423.

20. Chaput, D. L.; Bass, D.; Alam, M. M.; Hasan, N. A.; Stentiford, G. D.; Aerle, R. v.; Moore, K.; Bignell, J. P.; Haque, M. M.; Tyler, C. R., The segment matters: probable reassortment of tilapia lake virus (TiLV) complicates phylogenetic analysis and inference of geographical origin of new isolate from Bangladesh. Viruses 2020, 12, (3), 258.

