## Supplementary material for "Meta-transcriptomic identification of divergent *Amnoonviridae* in fish": Figure S1

**Figure S1.** Agarose gels electrophoresis showing PCR products from three sets of primers that target a region in the PB1 gene segment (RdRp) for ten individuals of *Meuschenia flavolineata*.

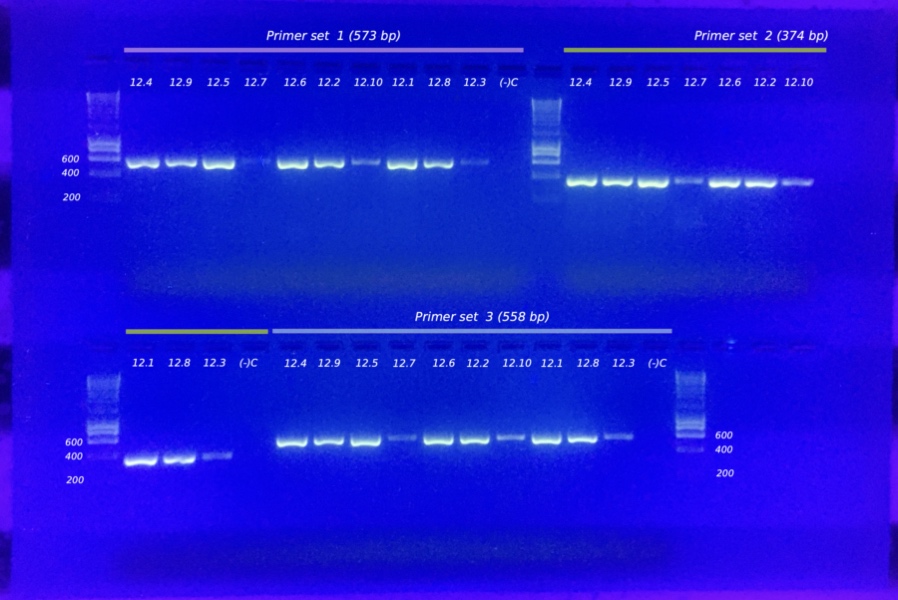
