## Supplementary material for "Meta-transcriptomic identification of divergent *Amnoonviridae* in fish": Table S1

**Table S1**. List of primer sets used for the RT-PCR confirmation of flavolineata virus in specimens of *Meuschenia flavolineata*.

| **Primer set** | **Primer ID (Forward/Reverse)** | **Sequence**  **(5’-3’)** | **Annealing temperature (ºC)** | **Target size (bp)** |
| --- | --- | --- | --- | --- |
| First | F1A-6F | CTTTCTGTTGGGCCCAGGAT | 64.8 | 573 |
|  | F1B-578R | GTTGAGCAGCGAACAAGTGG |  |  |
| Second | F3A-886F | GTGGAGTATCGGTTAGAGGAGATG | 64.6 | 374 |
|  | F3BA-1259R | TTCAGCACAGTCTTCCCACC |  |  |
| Third | F3A-886F | GTGGAGTATCGGTTAGAGGAGATG | 64.6 | 558 |
|  | F3B-1443R | AGGAGTCGAGGAGTTGGTGA |  |  |
