## Supplementary material for "Meta-transcriptomic identification of divergent *Amnoonviridae* in fish": Table S2

**Table S2**. All virus transcripts identified in this study that fell across genomic segments within the *Amnoonviridae*.

| **Virus species** | **Genomic segments (length, nt)** | **TSA accessions (if applicable)** |
| --- | --- | --- |
| *Flavolineata virus* | Segment 1 (1536 nt) | N/A |
|  | Segment 3 (1314 nt) | N/A |
|  | Segment 4 (1179 nt) | N/A |
| *Piscibus virus* | Segment 1 (270 nt) | N/A |
| *Dolomieu virus* | Segment 1 (1440 nt) | GDQU01066121.1, GDQU01106321.1, GDQU01283605.1, GDQU01532168.1 |
| *Namensis virus* | Segment 1 (1503 nt) | GHYH01080462.1, GHYH01005036.1, GHYH01084204.1 |
|  | Segment 2 (1254 nt) | GHYH01005036.1 |
|  | Segment 4 (900 nt) | GHYH01084204.1 |
| *Hamatus virus* | Segment 1 (321 nt) | GFMN01088333.1 |
| *Stewartii virus* | Segment 1 (1743 nt) | GIBO01031171.1,  GIBO01013027.1 |
|  | Segment 2 (1386 nt) | GIBO01013027.1 |
| *Plagiostomus virus* | Segment 1 (366 nt) | GHXZ01024367.1, GHXZ01079240.1 |
|  | Segment 2 (321 nt) | GHXZ01079240.1 |
| *Przewalskii virus* | Segment 1 (1761 nt) | GHYJ01002273.1,  GHYJ01008047.1,  GHYJ01010906.1 |
|  | Segment 2 (1350 nt) | GHYJ01010906.1 |
| *Asotus virus 1* | Segment 1 (1710 nt) | GHGF01026383.1, GHGF01034639.1, GHGF01033499.1, GHGF01028660.1, GHGF01037407.1 |
|  | Segment 2 (1362 nt) | GHGF01027066.1 |
|  | Segment 3 (1389 nt) | GHGF01033499.1 |
|  | Segment 4 (942 nt) | GHGF01028660.1 |
| *Asotus virus 2* | Segment 1 (1719 nt) | GHGF01016319.1, GHGF01027066.1, GHGF01047620.1 |
|  | Segment 2 (1368 nt) | GHGF01034639.1 |
| *Nudifrons virus* | Segment 4 (1032 nt) | HACN01008153.1 |
